# Multi-Level DBSCAN: A Hierarchical Density-Based Clustering Method for Analyzing Molecular Dynamics Simulation Trajectories

**DOI:** 10.1101/2021.06.09.447666

**Authors:** Song Liu, Siqin Cao, Michael Suarez, Eshani C. Goonetillek, Xuhui Huang

## Abstract

Molecular Dynamic (MD) simulations have been extensively used as a powerful tool to investigate dynamics of biological molecules in recent decades. Generally, MD simulations generate high-dimensional data that is very hard to visualize and comprehend. As a result, clustering algorithms have been commonly used to reduce the dimensionality of MD data with the key benefit being their ability to reduce the dimensionality of MD data without prior knowledge of structural details or dynamic mechanisms. In this paper, we propose a new algorithm, the Multi-Level Density-Based Spatial Clustering of Applications with Noise (ML-DBSCAN), which combines the clustering results at different resolution of density levels to obtain the hierarchical structure of the free energy landscape and the metastable state assignment. At relatively low resolutions, the ML-DBSCAN can efficiently detect high population regions that contain all metastable states, while at higher resolutions, the ML-DBSCAN can find all metastable states and structural details of the free energy landscape. We demonstrate the powerfulness of the ML-DBSCAN in generating metastable states with a particle moving in a Mexican hat-like potential, and four peptide and protein examples are used to demonstrate how hierarchical structures of free energy landscapes can be found. Furthermore, we developed a GPU implementation of the ML-DBSCAN, which allows the algorithm to handle larger MD datasets and be up to two orders of magnitude faster than the CPU implementation. We demonstrate the power of the ML-DBSCAN on MD simulation datasets of five systems: a 2D-potential, alanine dipeptide, β-hairpin Tryptophan Zipper 2 (Trpzip2), Human Islet Amyloid Polypeptide (hIAPP), and Maltose Binding Protein (MBP). Our code is available at https://github.com/liusong299/ML-DBSCAN.

## 1. Introduction

Molecular Dynamics (MD) simulations can reveal structural and dynamic details of molecular systems by providing spatial (up to atomic level) and temporal (up to femtosecond) information of complex biological macromolecules^1–3^. In recent years, the development of high-performance computing facilities has made long time MD simulations possible, allowing biological processes (of micro- or even milli-second timescales) to be simulated. However, MD simulations can yield up to a million different conformations, giving rise to enormous challenges that can hinder the understanding of the biomolecular processes being investigated.

Clustering algorithms^4–10^ have been developed and employed in MD simulations because of their ability to reduce the dimensionality of the trajectories produced by MD simulations. They then generate models that consist of fewer states by grouping conformations with similar geometry and structure; this produces a dataset that is much easier to understand and interpret. The most popular clustering algorithms are the k-centers, k-means, k-medoids, and density-based clustering algorithms, where k-centers^11^ clustering aims to minimize the maximum radius of all clusters; the k-means and k-medoids^12^ algorithms attempt to reduce the total distance of each conformation to its assigned cluster center, and the density-based clustering algorithms^13^ aim to find high population regions where metastable states or free energy minima may be situated.

These clustering algorithms are often used to generate conformational states in Markov State Models (MSMs)^14–27^ to elucidate biomolecular dynamics. The limitation of most existing clustering algorithms, however, is their inability to be used directly to identify kinetically metastable states, as they are based on geometric similarity between pairs of protein conformations. Consequently, a two-step scheme is often adopted in MSMs to determine the metastable states^14,28^; the first step clusters or splits the conformational space into microstates based on geometric similarity of protein conformations, and the second step identifies the metastable states by lumping these microstates based on their kinetic proximity. However, this two-step scheme can only provide limited information about the free energy landscape, as the hierarchical structures of metastable states are excluded. Therefore, improved clustering algorithms are required to generate metastable states across different timescales or to study the hierarchical structure of the free energy landscape.

Density-based clustering algorithms are particularly suitable for the study of biomolecules as they can be used to directly determine the metastable states of proteins^29^ that are situated at free energy basins (high density regions). In particular, the density-based clustering algorithms are shown to be effective to resolve conformational states that contain geometrically similar conformations but are separated by kinetic barriers^13,30,31^. These advantages have led to the specific development of different density-based clustering algorithms for the study of biomolecular dynamics; these include the neighbor-based algorithms^32^, density-peaks^33^, Robust-Distance Based, and the Adaptive Partitioning by Local Density-peaks (APLoD)^13^ algorithm.

The Density-Based Spatial Clustering of Applications with Noise (DBSCAN) algorithm^34^ is one of the most popular density-based clustering methods, and has been widely used in analyzing MD datasets; notably in core-set MSM that utilizes this algorithm to find high population regions^35^. The DBSCAN works by adopting the local density, defined as the number of neighboring points, to find high-density or metastable states. When using the DBSCAN, MD conformations at high-density regions are grouped into clusters, and outliers in low-density regions are regarded as “noise”. The density-based clustering algorithms can efficiently elucidate the features of the free energy landscapes underlining biomolecular dynamics, where the metastable conformational states (free energy minima) are highly populated and thus identified by density-based clustering algorithms. Therefore, the DBSCAN holds great promise to be applied to analyze MD simulations of biomolecules^9^.

Although density-based clustering methods show success in analyzing biomolecular dynamics, DBSCAN in particular performs clustering according to a single and uniform threshold to define the high-density regions. However, the free energy landscapes of biomolecules contain a hierarchical architecture with free energy minima separated by free energy barriers at different density levels^36–38^. Different timescales can be revealed by changing the resolution: e.g. large-scale folding or unfolding of the protein backbone correspond to deep free energy minimum separated by substantial free energy barriers, and thus occurs at slow timescales. On the contrary, sidechain rotations and other local conformational changes of proteins should describe faster transitions between shallower free energy minimum. As the hierarchical structure of the free energy landscape is particularly important for understanding biological processes and functions of biomolecules, new algorithms are required to reveal this hierarchical structure. We have previously attempted to reveal the hierarchical structure of a free energy landscape by developing the Hierarchical Nyström method^38^, which can effectively treat both the highly populated and sparsely populated regions to finally obtain dominant metastable states. However, this method still requires kinetic lumping at different density resolution levels. We noticed that the DBSCAN algorithm is particularly suitable for the hierarchical analysis of the free energy landscape, as the alteration of two parameters (the radius for finding the neighboring points, and the minimal number in the neighborhood) allows the algorithm to be performed at different resolutions of the free energy landscape. The combination of these clusters at different resolutions can be used to reveal the hierarchical structure of the conformational space as well as the dynamics of large biomolecules. This motivated us to develop a multi-level DBSCAN (ML-DBSCAN) clustering algorithm in this study.

In this paper, we propose a ML-DBSCAN clustering algorithm that determines the hierarchical structure of the free energy landscape of biomolecules by combining DBSCANs performed at different resolutions. The ML-DBSCAN can show the metastable states at each resolution level, generating a hierarchical structure of the free energy landscape similar to one obtained through the Nyström method. The state assignment of all resolution levels can be combined into a metastable state assignment that relates to the free energy minima. We have developed two implementations of the ML-DBSCAN; one for the CPU and the other one for GPU. We also present five examples to demonstrate both the workflow and powerfulness of our algorithm and implementations. We anticipate that our method will be widely applicable to biomolecular systems. The organization of this manuscript is as follows: we first introduce the algorithm of the DBSCAN and the ML-DBSCAN. Then, we present our GPU implementation of our ML-DBSCAN. Finally, we demonstrate the power of the ML-DBSCAN on MD simulation datasets of five systems: a 2D-potential, alanine dipeptide, β-hairpin Tryptophan Zipper 2 (Trpzip2), Human Islet Amyloid Polypeptide (hIAPP), and Maltose Binding Protein (MBP).

## 2. Methods

### 2.1. The DBSCAN Algorithm

The DBSCAN algorithm is a density-based clustering method that can effectively obtain metastable high-density states from massive MD conformations. In this algorithm, metastable states are defined as high-density regions, given by the large enough number of neighbors (these neighbors are identified in a low dimensional space, see Sec 2.2. for details). Therefore, MD conformations in low-density regions shall not exert sufficient number of neighbors and thus are regarded as outliers in the DBSCAN algorithm. One of the parameters that control the DBSCAN algorithm, *MinPts*, represents the minimum number of conformations in the given neighborhood (*N(p)*) and is used to identify outliers. The DBSCAN algorithm can use this parameter to define the minimal density (i.e., number of neighboring conformations) needed to prevent the conformations from being defined as outliers and excluded in the subsequently analysis. The second parameter that controls the algorithm, *ε*, represents the maximum distance between neighbors and is used to define the radius of the neighborhood.

The two parameters of the DBSCAN, *ε* and *MinPts*, can be used to control the resolution, allowing the DBSCAN to perform multi-resolution analysis. When *ε* is large and *MinPts* is small, the whole MD dataset may be grouped into one state; but when *ε* is small and *MinPts* is large, a larger number of states will be generated and more details of the free energy landscape are found. Therefore, by tuning (*ε, *MinPts**), the DBSCAN can be performed at multiple resolutions.

Our implementation of the DBSCAN consists of four steps: the neighborhood search, the generation of new clusters, cluster growth, and outlier treatment. In the first step, all the neighbors within the radius *ε* of each conformation are found, and core conformations are identified. We define a core conformation as having at least *MinPts* conformations within its distance *ε*. In the second step, new clusters are generated at each core conformation if the conformation hasn’t been assigned to any cluster. In the third step, all the density connected conformations are assigned to the same cluster as the core conformations; and in the fourth step, the remaining unclassified conformations in the dataset are labeled as outliers.

### 2.2 Multi-Level DBSCAN Algorithm

The DBSCAN algorithm introduced in Sec. 2.1 can only perform clustering at a single density level, i.e., a single and uniform definition of high-density regions determined by. In this section, we introduce a Multi-Level DBSCAN (ML-DBSCAN) method that performs the DBSCAN clustering at different density levels and further combines the results to obtain a hieratical set of clusters that reflect the structure of the underlying free energy landscape. A simple 1-D free energy example suited for ML-DBSCAN is displayed in Fig. 1 A, and this 1-D free energy profile contains energy minima located at different density levels.

**Figure 1:**
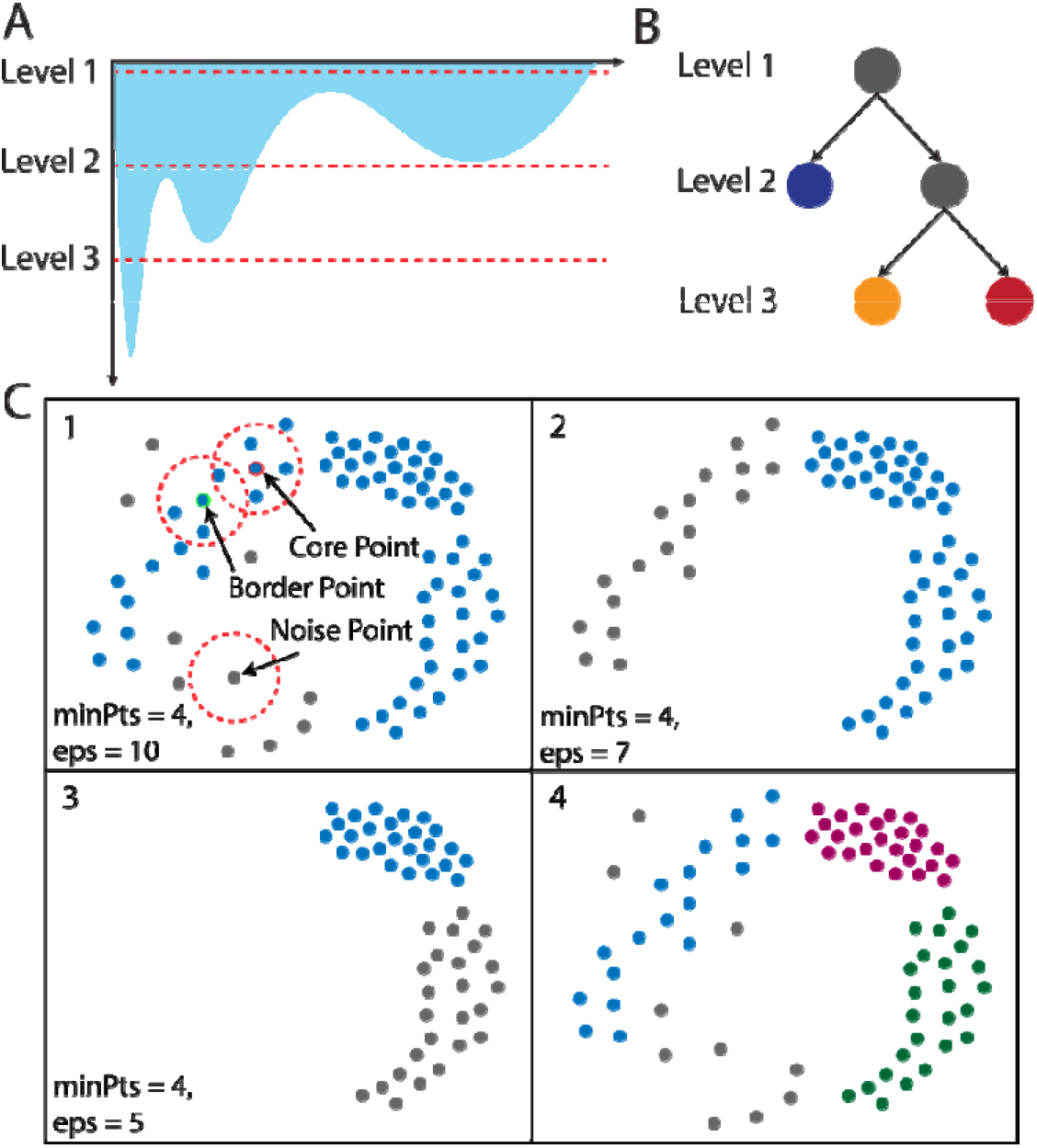
Schematic diagram for the ML-DBSCAN algorithm. (A) 1D free energy landscape. (B) Hierarchical clustering of Multi-level DBSCAN on a 1D free energy landscape. (C) The ML-DBSCAN clustering algorithm on a synthetic 2D dataset. The grey points are regarded as “noise”. The point with a red outline within the red dashed circle of the radius of eps is a core point, and the point with the green outline is a border point.

Our ML-DBSCAN was implemented in the following way: we first partitioned the MD conformations with various densities by setting up a group of DBSCANs that ran at different resolutions. The resolutions were arranged by a different combination of parameters before running the ML-DBSCAN, which works as follows: First, dense areas are separated using a DBSCAN instance with parameters favoring high density data points, discarding remaining points as outliers. Then, the sparse density areas can be separated by selecting another DBSCAN instance with parameters favoring low data point density, additionally revealing clusters among the previously discarded points. After the DBSCAN is performed at various resolutions, a hierarchical result can be obtained by combining the DBSCANs at different resolution levels, as shown in Fig. 1.

The implementation of our ML-DBSCAN involves four parts, as shown in Fig. 2. In part a, “MD trajectories”, we first define a space that defines the L2 distances between conformations. This step is achieved by featurization of MD data (e.g., using tICs from the time-structure based Independent Component Analysis (tICA)^39,40^). In part b, we use automatic hyper-parameter optimization to determine the upper and lower bound of both *ε* and *MinPts*. In this part, we perform some test DBSCANs to estimate the regions of (*ε, *MinPts**), and then generate a number of (*ε, *MinPts**) in this range. In part c, “Multi-level DBSCAN”, DBSCANs at the (*ε, *MinPts**) obtained in the previous part are performed. This step is performed in a two-stage manner, where in the first stage, low-resolution DBSCANs will be performed, which generates only one metastable state containing all dense regions of the free energy landscape. In this stage, the outliers are found and discarded to allow us to focus on the dense regions in subsequent steps. In the second stage, a number of DBSCANs at higher resolutions will be performed to uncover more details of each metastable state; this stage is achieved by further splitting the large metastable states into a number of small states. Also, since each DBSCAN is run independently, this stage can be performed in parallel. Finally, in part d, “hierarchical assignment”, the hierarchical information of all resolution levels is collected, where both the state assignment spanning a wide range of resolutions and the hierarchical structure of free energy landscape can be obtained.

**Figure 2:**
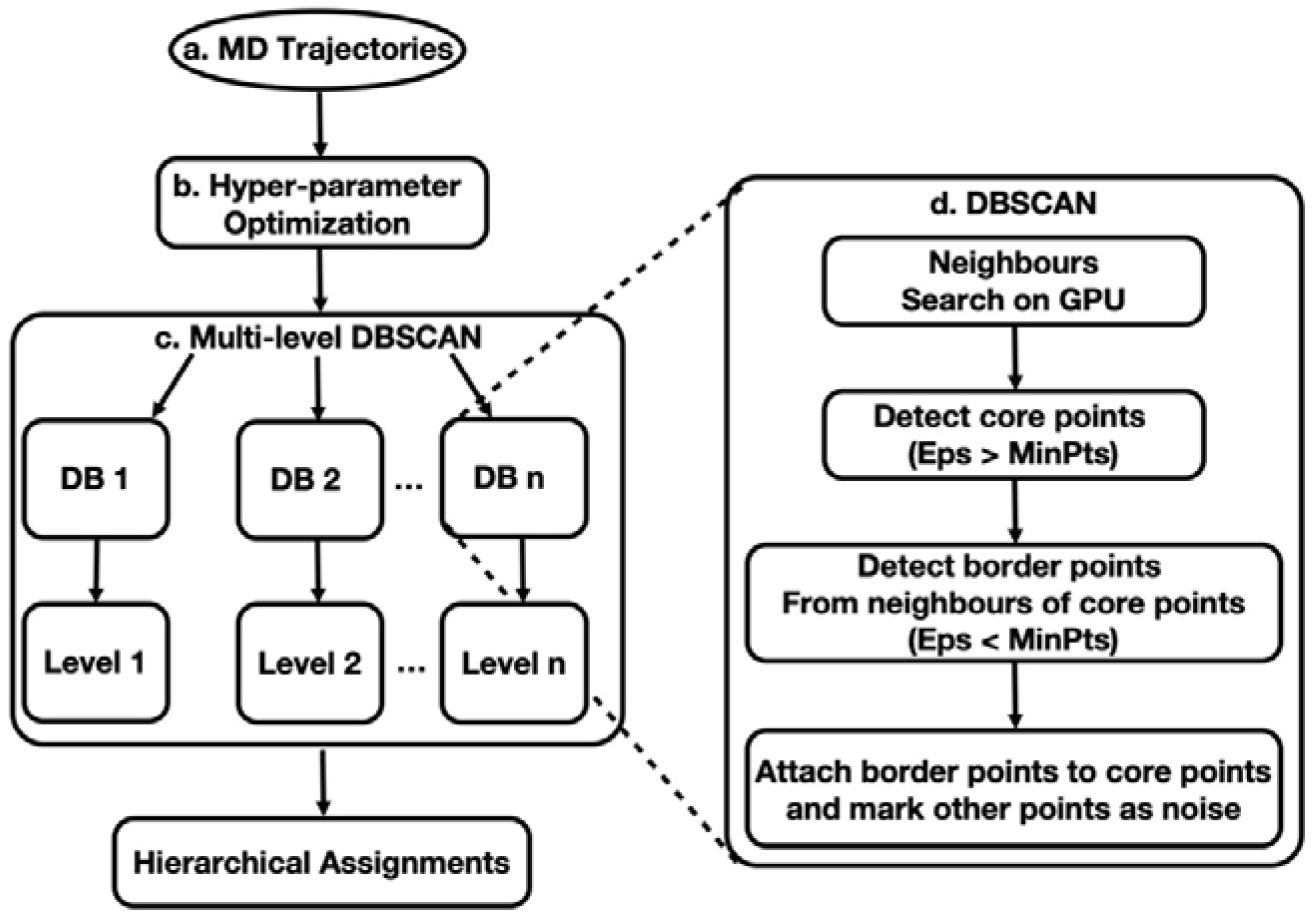
Flowchart of the ML-DBSCAN algorithm.

**Figure 3.**
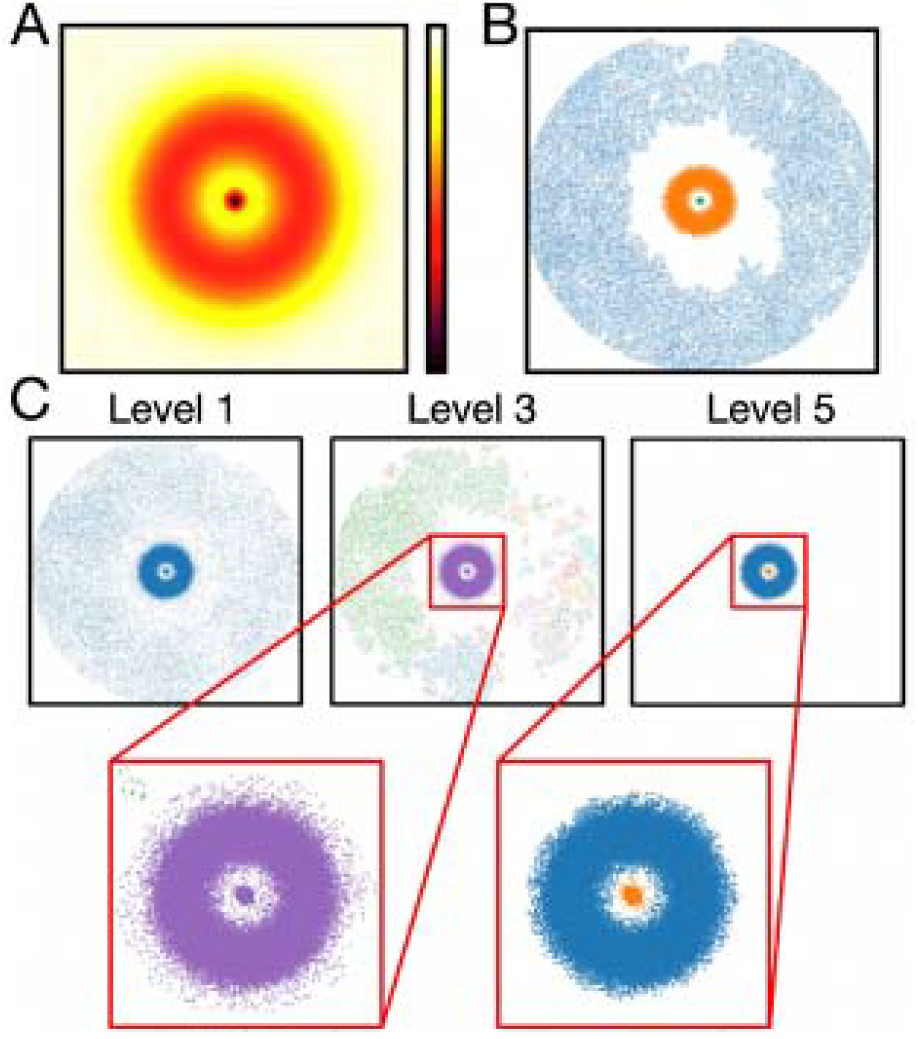
The ML-DBSCAN successfully identifies core states on the “Mexican Hat” potential dataset. The needle-like central minimum (A) mimics a multi-body protein-ligand binding process. The central needle-like minimum, the ring-like intermediate minimum (red), and the outer broad and shallow minimum (yellow), represent the bound form of the protein-ligand complex, the state in which the ligand first encounters the protein, and the diffusion state where the ligand diffuse freely in solution without any contact to the protein, respectively. (B) Identified clusters on the finite sampling of this potential (C) Identified large metastable clusters of each level with a population greater than 1.0%.

The protocol of the ML-DBSCAN is demonstrated in Fig. 1B & C using a one-dimensional free energy landscape as shown in Fig. 1A. This free energy landscape has a deep minimum that can always be identified as cluster by all levels of the free energy landscape, and two other minima that can be found at level 2 and 3, respectively. When performing a ML-DBSCAN, we first find level 1, which consists of a single metastable state that covers all data regions. Then, at level 2, the huge metastable state at level 1 will split into two metastable states that is separated by the highest energy barrier. Finally, at level 3, the metastable state at level 2 further splits into two states. When building the ML-DBSCAN, a hierarchical family tree of metastable states at different levels is automatically built (Fig. 1B), where each parent node corresponds to a large dense region that contains conformations of all child nodes. In addition, the family tree displayed in Fig. 1B reveals the hierarchical structure of the free energy landscape, and a metastable state assignment (Panel 4 in Fig.1C) spanning all the three free energy levels (level 1-3 in Fig. 1A) can be obtained by combining the state assignment as shown in Panels 1-3 in Fig. 1C.

### 2.3 The GPU implementation of the Multi-Level DBSCAN Algorithm

The ML-DBSCAN is sufficiently efficient for simple datasets (with a small number of MD conformations) such as those shown in Fig. 1, 4 and 5. However, for larger dataset (e.g., millions of MD conformations, see Fig. 6), the efficiency of the ML-DBSCAN decreases significantly. In order to analyze these larger datasets, a highly efficient implementation of the ML-DBSCAN is required. Since the ML-DBSCAN is highly parallelizable (as demonstrated in Sec. 2.2 and Fig. 2), a GPU implementation can be adopted to improve the efficiency.

**Figure 4.**
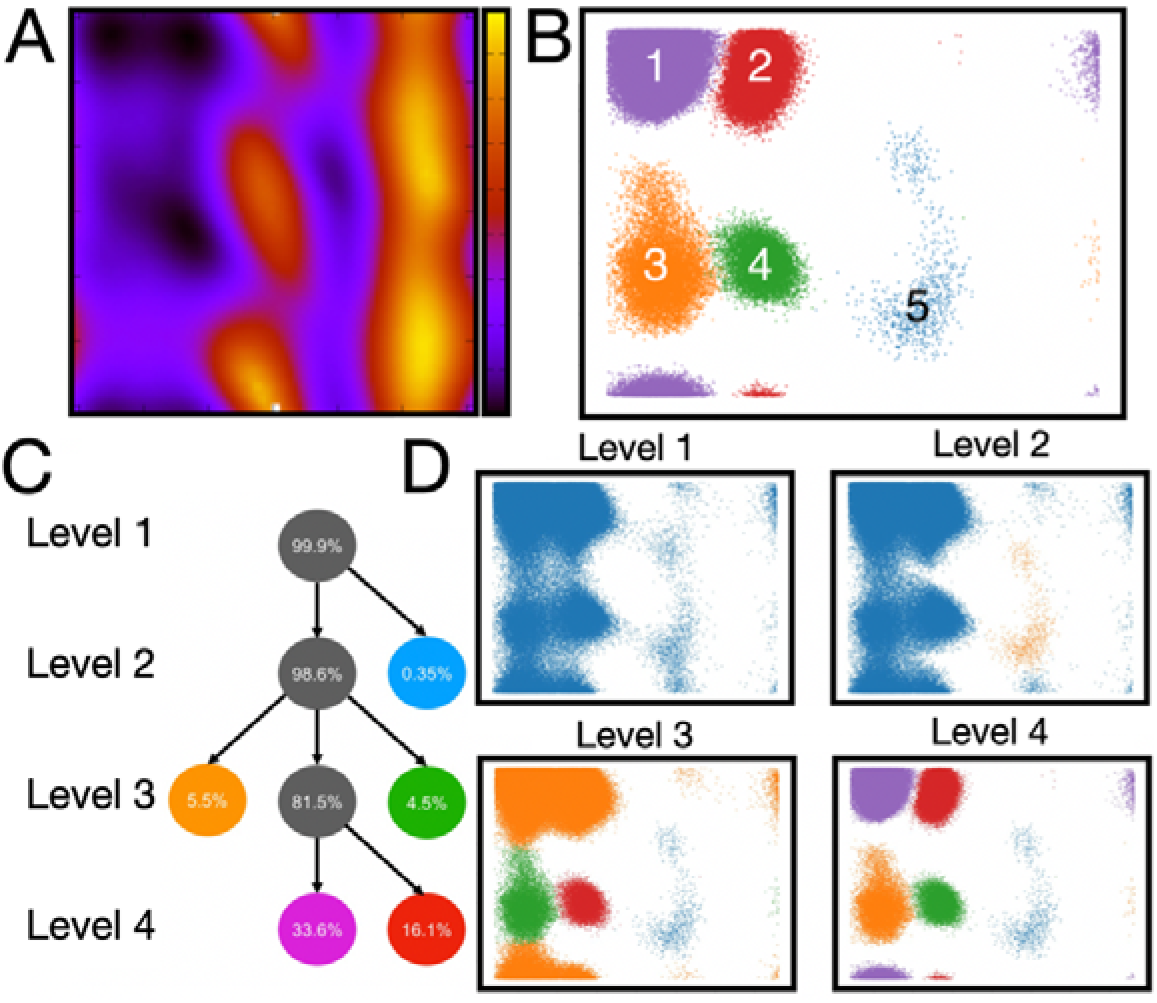
A four-level ML-DBSCAN analysis identifies five free energy minima of the Alanine Dipeptide dataset. (A) The density of Alanine Dipeptide conformational landscape (B) Identified core states clusters by ML-DBSCAN (C) A hieratical set of density levels identified by ML-DBSCAN (D) Identified the large metastable clusters at each level.

**Figure 5.**
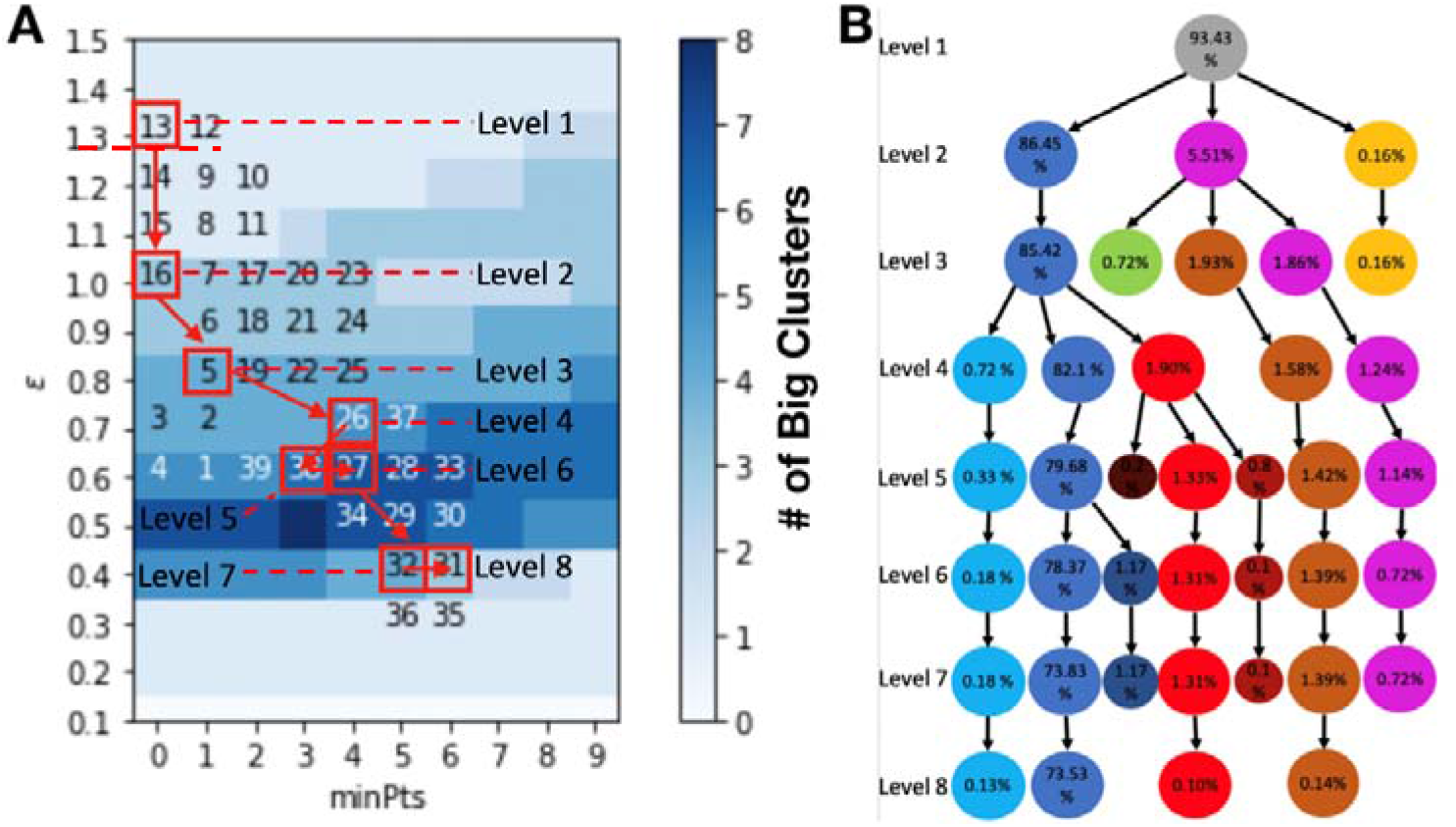
A Monte-Carlo search for the hyperparameters in the ML-DBSCAN analysis of the Trpzip2 peptide folding. (A) Exploring for a better set of DBSCAN hyper-parameters of each level based on the Monto Carlo search. (B) The hierarchical structure of the ML-DBSCAN, where each node represents a metastable state.

**Figure 6.**
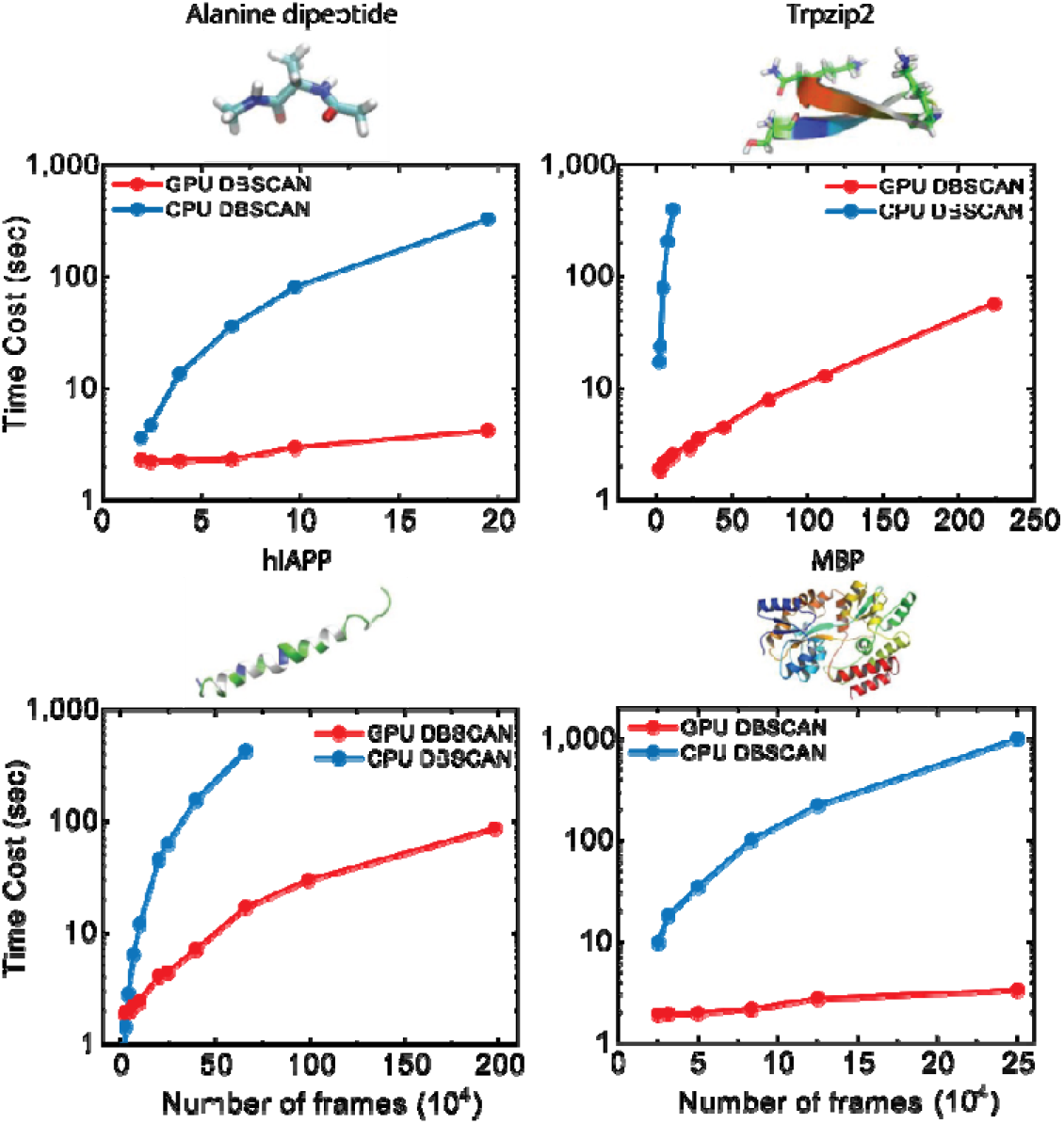
The GPU implementation of the DBSCAN algorithm can achieve acceleration up to two orders of magnitude when comparing to the CPU implementation. Four systems are evaluated: the alanine dipeptide, 12-residue -hirpin Tryptophan Zipper 2 (TripZip2), Human Islet Amyloid Polypeptide (hIAPP), and MaltoselJbinding protein (MBP). Time costs are compared between the two clustering algorithms; blue curves are the results of CPU implementations and red curves are the results of GPU implementations. The CPU implementations are run on Intel(R) Xeon(R) CPU E5-2680 v4 @ 2.40GHz, while the GPU implementations are run on a NVIDIA GeForce GTX 1080 Ti.

We have developed a GPU implementation of the ML-DBSCAN that involves running multiple DBSCANs in parallel, as well as parallelizing the DBSCAN algorithm. The parallelization of the DBSCAN algorithm is achieved by FAISS, a Python library readily available on GPU that can significantly accelerate the nearest neighbor search engine^35^. To further improve the speed of the DBSCAN, two high-speed methods of FAISS have been used in our ML-DBSCAN for the nearest neighbor search: an approximate nearest neighbor (ANN)^41^ algorithm for Approximate Similarity Searches, and IndexIVFFlat for inverted indexing (IVF)^42^. For the Approximate Similarity Searches, we use the more time efficient ANN instead of the more rigorous k-nearest neighbor (KNN) algorithm^32^. Meanwhile, we use IndexIVFFlat for IVF as it is the most efficient algorithm for the nearest neighbor search.

### 2.4 Simulation Datasets

To demonstrate the power of our ML-DBSCAN algorithm, we have applied it to five MD simulation datasets.

#### 2D-potential

The first one is a “Mexican hat” like 2D-potential^30^ that is used to demonstrate the performance of the ML-DBSCAN as it mimics a multi-body protein-ligand binding process. Three energy minima can be found on the potential energy surface, with the central minima much deeper and narrower than the other two minima (see Fig. 5A). As a result, the dynamics in the needle-like central minimum are significantly slower than other minima. 100,000 conformations sampled by MD simulations of this system taken from our previous study^30^ are used for our ML-DBSCAN analysis.

#### Four Biological Systems

The four other datasets are MD simulations of peptides and proteins. The first MD dataset is the dynamics of a terminally blocked alanine peptide, Ac-ala-NHMe, that was generated in explicit solvents by Chodera *et al*^21^; this dataset contains 975 trajectories and 195,000 conformations in total. The second dataset is the folding dynamics of a 12-residue Trpzip2 peptide^43^; this dataset contains 2,239,523 MD conformations. The third dataset contains MD simulations of a 37-residue intrinsically disordered hIAPP peptide; this dataset contains 20 trajectories at a length of 198-ns and 1,980,000 conformations in total^44^. The fourth dataset is the 370-residue MBP protein^11^; this dataset consists of 25 trajectories, each performed for 50 ns with a saving interval of 5-ps and 0.25 million conformations in total. For each of these four datasets, we first use pairwise distances of all heavy atoms to perform (tICA)^39,40^. Four tICs are chosen from each dataset at a tICA lagtime of 10 ps, 1 ns, 20 ns, and 50 ns for the Alanine dipeptide, Trpzip2, hIAPP and MBP, respectively. The L2 distances of a four-dimensional space consisting of the first four tICs are used as distances between data points in the DBSCANs.

## 3. Results and Discussions

In this section, we use the five examples mentioned above to demonstrate the workflow and power of our ML-DBSCAN algorithm. First, we show that the ML-DBSCAN is applicable for a highly heterogeneous free energy landscape, like the one underlying the Mexican hat potential (see Sec. 3.1). Second, we use an alanine dipeptide to demonstrate the power of the ML-DBSCAN to obtain the hierarchical structure of the free energy landscape of large biomolecules in Sec. 3.2. Third, we demonstrate the construction of the ML-DBSCAN using the Trpzip2 dataset in Sec. 3.3. Finally, we show improved efficiency with the GPU implementation of the DBSCAN with the MD trajectories of Trpzip2, hIAPP, and MBP in Sec. 3.4.

### 3.1 ML-DBSCAN reveals heterogeneous energy minima in a 2D-potential

We first demonstrate how the ML-DBSCAN can identify multiple energy minima with drastically different depth in the “Mexican hat” 2D-potential. The “Mexican hat” potential has three heterogeneous energy minima as shown in Fig. 3A, where three metastable states (the yellow and black regions) can be found at each energy minimum: the origin and the two broad ring-like regions. This “Mexican hat” potential is an analog to many protein-ligand systems, where the broad ring-like region represents the free diffusion of the ligand, the narrow ring line represents the first encounter that the protein has with the ligand, and the minimum at the origin represents the bound states. Previous studies^30^ show that this “Mexican hat” potential impose challenges for clustering methods, including k-centers^11^ and k-means^45^. In MSMs, an additional step called “lumping” is required to get the metastable state assignments, while in the DBSCAN, the “Mexican Hat” potential can be properly handled to obtain all the metastable states without having to perform “lumping”. Fig 3B shows that the three metastable states (labelled in blue, yellow, and dark blue) can be clearly obtained from the DBSCAN (Fig. 3B is established by combining the results obtained from resolution level 1, 3, and 5, which are shown in Fig. 3C). The “Mexican hat” example can also show how the DBSCAN can be used to obtain the potential landscape at different resolutions. As shown in Fig. 3C, the DBSCAN with the lowest resolution can be performed at level 1, which has only one state. At level 3, the DBSCAN can separate the joint middle ring-like region and the needle-like region as another big state; and at level 5, the middle ring-like region and the needle-like region can be distinguished from each other.

The details of performing the ML-DBSCAN for the “Mexican hat” potential are shown in Fig. S1. In the ML-DBSCAN, we first defined a parameter grid in the range of *ε* and *MinPts* at *ε* E [1, 3] and *MinPts* ∈ [15, 20]. Then, we performed 25 DBSCANs at *ε* = 1, 1.5, 2, 2.5, 3 and *MinPts* = 15, 16, 18, 19, 20 for every combination of the two parameters. We noticed that the DBSCANs of the upper-right triangle region of the (*ε, *MinPts**) space (large *ε* and *MinPts*) overwhelmingly shows one big cluster in the sparse region, while the lower-left triangle region (small *ε* and *MinPts*) tends to remove the sparse region as noise and splits the densest needle region with the middle ring-like region as different clusters. Along the diagonal of the parameter grid the DBSCANs performed at those specific (*ε, *MinPts**) show the best partitioning and are therefore used for further analysis and graphing.

### 3.2 Hierarchical analysis of the free energy landscape of Alanine Dipeptide by the ML-DBSCAN

In this section, we use the alanine dipeptide as an example to demonstrate how to find the hierarchical structure on the free energy landscapes of peptide systems using the ML-DBSCAN. The projection of the free energy landscape onto a pair of torsional angles of the alanine dipeptide is shown in Fig. 4A. As shown in Fig. 4B, five free energy minima (state 1, the C5 state; state 2, the PII state; state 3, *α_P_*; state 4, *α_R_*; and state 5, 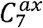) can be successfully found by our ML-DBSCAN.

The hierarchical analysis of this free energy landscape enabled by our ML-DBSCAN analysis is displayed in Fig. 4C: choosing the parameters for level 1 with the lowest resolution leads to all the conformations being clustered to one state; while tuning the hyper-parameters to higher resolutions, level 2, 3, and 4, reveal the blue, yellow/green, and purple/red states, respectively. State C5 and PII are separated with a barrier of 1 kcal/mol. Therefore, the splitting of state C5 and PII can only be seen with the highest resolution of the DBSCAN, level 4. State C5/PII and *α_P_/α_R_* are separated by a barrier of 2 kcal/mol, and can be seen at the resolution of level 3. In addition, a high energy barrier of 6~7 kcal/mol between state C5/PII/*α_P_/α_R_* and 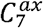 can be seen in the first resolution level, level 1. This example clearly shows that different metastable states corresponding to different free energy minima would appear at different resolutions, so a multi-level approach (like the ML-DBSCAN) that scans through all the resolutions is required to obtain all the metastable states.

The protocol for finding the hierarchical structure of the free energy landscape is further shown in Fig. 4C-D and Fig. S2. We first chose a wide range of *ε* and *MinPts* that could cover every resolution level. Then, we found that the DBSCANs performed at the diagonal of the whole range of (*ε, *MinPts**) can be used as different resolution levels in the ML-DBSCAN (see Level 1 to Level 8 in Fig. S2). At (*ε, *MinPts**) = (0.08, 5) at top-left region of Fig. S2, the whole dataset is grouped into one single state, and this state assignment is called Level 1. Furthermore, at (*ε, *MinPts**) = (0.06, 15), which is chosen as level 2, the whole dataset splits into two states, the 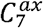 state and a big state that contains all the other four metastable states. At a higher resolution (Level 3), (*ε, *MinPts**) = (0.03, 30), the large state at level 2 (blue in Fig. 4D, Panel Level 2) will further split into three states: aP, aR, and another big state that consists of all beta structures. Finally, at Level 4, the bottom-right corner of Fig. S2, (*ε, *MinPts**) = (0.01,40), the beta structures state will further split into C5 and PII, and thus all five states have been found.

### 3.3 ML-DBSCAN can reveal the hierarchical free energy landscape of Trpzip2

In this section, we use a large biomolecule, Trpzip2, to demonstrate the efficiency of our algorithm in search for resolution levels that can quickly construct the hierarchical structures of the free energy landscape. The Trpzip2 folding has been widely studied through both experiments and MD simulations^46^. In particular, the metastable states during its folding process have been revealed by MSMs built from extensive MD simulations^11,21,47^. Since the Trpzip2 is a 12-residue peptide, with substantially larger size than the alanine dipeptide, and a systematic search of the space in ML-DBSCAN at comparable detail to the alanine dipeptide example would require a lengthy computation time. Therefore, a more efficient method to search for resolution levels is needed for large peptide systems like Trpzip2.

In order to find proper resolution levels, we developed an automatic hyper-parameter optimization method that utilizes a Monto Carlo search of the hyper-parameter space rather than a simple and exhaustive Grid Search (see Fig. 5A). In our approach, we adopt a random walk scheme to search for the efficient regions among all the resolution levels. Firstly, is randomly initialized as a starting point on a pre-defined grid-space and the DBSCAN is computed. Secondly, a step is taken on this grid-space in a random direction (up, down, left, right) and the DBSCAN is computed for this parameter set. If the random walk hits the boundary of parameter grid, then a prior non-walked direction is randomly continued as the new step. This is iteratively repeated until a defined maximum step parameter (n) is reached. This leads to n number of DBSCAN computations using n different combinations. Finally, we construct the hierarchical levels (5B) by eliminating the hyper-parameter combinations that show the same number of big clusters in the DBSCAN computation. Here, we select 8 levels as demonstrated by the red boxes, with each level leading to a different number of clusters at different resolution. In our random walk scheme, we only focus on large clusters which have more than 0.1% of the total population and ignore the outliers below this threshold. The hierarchical structure of the free energy landscape of Trpzip2 is shown in Fig. 5B.

### 3.4 GPU implementation greatly accelerated the ML-DBSCAN analysis of MD simulation datasets of larger peptide and protein systems

When applying the ML-DBSCAN to large simulation systems that typically contain millions of frames, a significant amount of computing time may be required for the ML-DBSCAN with the random walk scheme to find resolution levels (e.g., Fig. 6B). Therefore, in order to further improve the efficiency of the ML-DBSCAN, we developed the GPU implementation of the DBSCAN.

The GPU implementation of the DBSCAN can greatly increase its efficiency by two orders of magnitude for complex biomolecular systems. As shown in Fig. 6, the GPU implementation is one order of magnitude faster than the CPU implementation for small datasets (~ MD conformations) and can be two orders of magnitude faster for large datasets (~ MD conformations). Fig. 6 also shows the efficiencies of our GPU accelerated DBSCAN algorithm and the CPU implementation on four protein systems: the alanine dipeptide (2 residues, 200 k frames), Trpzip2 (12 residues, 2.5 million frames), hIAPP (37-residue, 2 million frames), and MBP (370 residues, 250 k frames) (see Fig. 6 for details) on Intel(R) Xeon(R) CPU E5-2680 v4 @ 2.40GHz and NVIDIA GeForce GTX 1080 Ti. Out of these examples, we find that when the ML-DBSCAN with the CPU implementation is applied to small datasets like hIAPP (Fig.6C) and MBP (Fig. 6D), which contain hundreds of thousands of frames, only ~ seconds of computing time is needed. However, larger datasets, that contains millions of frames, like Trpzip2, may require several hours of computing time. Therefore, as the number of frames increase from thousands to millions, the time required for our CPU implementation of Trpzip2 drastically increases from 20 seconds to several hours. Conversely, our GPU implementation of the DBSCAN can successfully analyze a Trpzip2 dataset with over two million frames within minutes. This shows that our GPU implementation of the DBSCAN is much more effective for ultra-large MD simulation datasets.

The density-based clustering algorithms are naturally suitable for the construction of core-set MSMs^48,49^. Our DBSCAN algorithm is not a full partitioning of the conformational space, as outlier MD conformations are discarded. Thus, metastable states generated by DBSCAN can serve as the cores of core-set MSMs, while the connection of these cores can be performed with other methods such as milestoning^50^. In the past, the DBSCAN could only provide results in one resolution, however, our ML-DBSCAN can provide metastable states in many resolutions. Therefore, we anticipate that our ML-DBSCAN algorithm may greatly assist the core-set MSM analysis of protein dynamics^15,19,25^, especially on the studies of functional conformational changes of biomolecules^51–56^.

## 4. Conclusion

In this study, we developed a ML-DBSCAN clustering algorithm to generate metastable states and hierarchical structures on free energy landscapes from large-scale MD trajectories. The ML-DBSCAN can be used to find metastable states on a highly heterogeneous dataset, which is challenging for other density-based and non-density-based clustering methods that only operate on one resolution. The combination of DBSCANs at different resolution levels can be used to find the hierarchical structure of the free energy landscape of large biomolecules. In addition, we developed a random walk scheme to rapidly search for resolution levels, and a GPU implementation of DBSCAN that improves the efficiency of the ML-DBSCAN by two orders of magnitude. With these developments, our ML-DBSCAN algorithm can be applied to ultra-large datasets containing millions of MD conformations. We anticipate that the ML-DBSCAN can facilitate the MSM analysis of biomolecular dynamics.

## 5. Acknowledgement

X.H. acknowledges the support from the Padma Harilela Endowment Fund.

## 6. Code Availability

The source code for ML-DBSCAN is available at https://github.com/liusong299/ML-DBSCAN.

