## Supplemental materials for "Multi-Level DBSCAN: A Hierarchical Density-Based Clustering Method for Analyzing Molecular Dynamics Simulation Trajectories"

### Supplementary Figures:

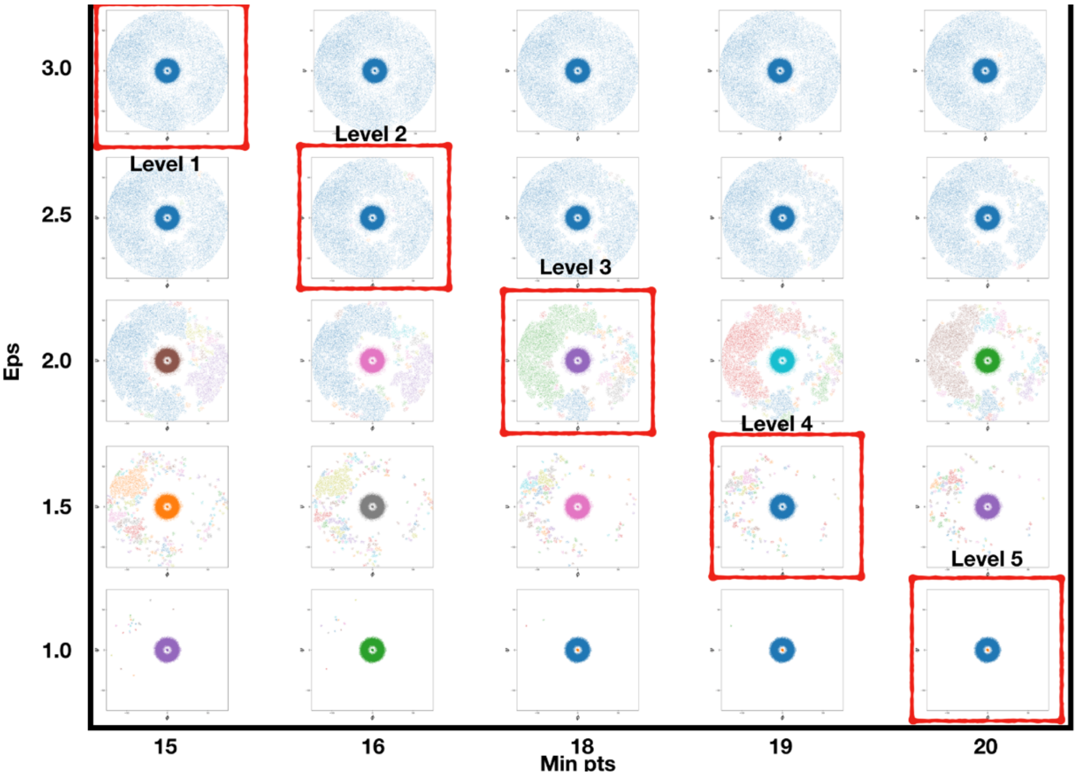


**SI Figure 1.** ML-DBSCAN identifies core states on the “Mexican hat” Potential dataset, especially the needle-like central minimum, parallel scanning of resolutions on Eps and Minpts, and level selection.

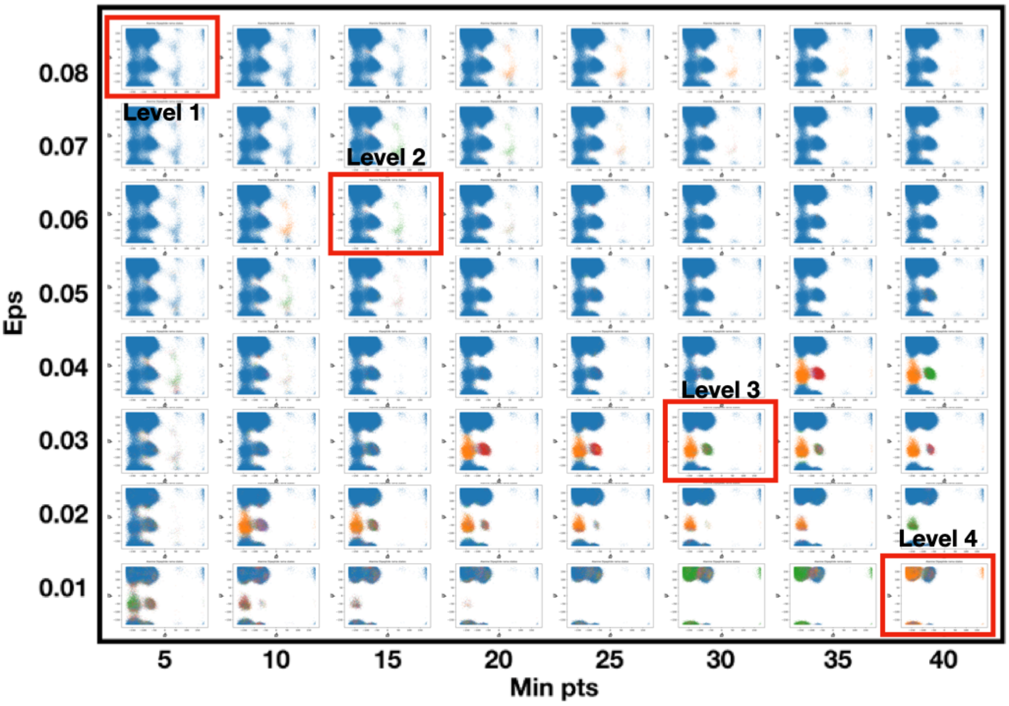


**SI Figure 2**. Parallel scanning of resolutions on Eps and Minpts, and level selection of the Alanine Dipeptide dataset
